# LDL-Binding IL-10 Reduces Vascular Inflammation in Atherosclerotic Mice

**DOI:** 10.1101/2024.03.04.582839

**Authors:** Lisa R. Volpatti, Salvador Norton de Matos, Gustavo Borjas, Joseph Reda, Elyse A. Watkins, Zhengjie Zhou, Mindy Nguyen, Ani Solanki, Yun Fang, Jeffrey A. Hubbell

## Abstract

Atherosclerosis is a chronic inflammatory disease associated with the accumulation of low-density lipoprotein (LDL) in arterial walls. Higher levels of the anti-inflammatory cytokine IL-10 in serum are correlated with reduced plaque burden. However, cytokine therapies have not translated well to the clinic, partially due to their rapid clearance and pleiotropic nature. Here, we engineered IL-10 to overcome these challenges by hitchhiking on LDL to atherosclerotic plaques. Specifically, we constructed fusion proteins in which one domain is IL-10 and the other is an antibody fragment (Fab) that binds to protein epitopes of LDL. In murine models of atherosclerosis, we show that systemically administered Fab-IL-10 constructs bind circulating LDL and traffic to atherosclerotic plaques. One such construct, 2D03-IL-10, significantly reduces aortic immune cell infiltration to levels comparable to healthy mice, whereas non-targeted IL-10 has no therapeutic effect. Mechanistically, we demonstrate that 2D03-IL-10 preferentially associates with foamy macrophages and reduces pro-inflammatory activation markers. This platform technology can be applied to a variety of therapeutics and shows promise as a potential targeted anti-inflammatory therapy in atherosclerosis.

## INTRODUCTION

The buildup of plaque in arterial walls, known as atherosclerosis, is a primary cause of coronary artery disease. Plaques are formed when cholesterol, lipids, cells, and extracellular matrix accumulate in the intima of arterial walls. Preventative measures and early interventions focus on reducing systemic cholesterol-laden low-density lipoprotein (LDL) through lifestyle changes or medications^1, 2^. While cholesterol-lowering statins are one of the most prescribed drugs, many patients still require surgical intervention at later stages of disease^3, 4^, and cardiovascular diseases remain the leading cause of death in the world^5, 6^.

Decades of research into the molecular mechanism of atherosclerosis has revealed that chronic inflammation is an important mediator of disease progression^7, 8, 9^. When macrophages or other cell types engulf excess lipids, they become pro-inflammatory foam cells that contribute to the plaque’s necrotic core^10, 11^. The CANTOS trial showed that a reduction of systemic inflammation is sufficient to significantly reduce cardiovascular events, independent of cholesterol lowering^12^. However, the anti-inflammatory therapy in this trial (canakinumab, a neutralizing monoclonal antibody against IL-1β) was associated with a higher incidence of fatal infections, suggesting the need for more targeted approaches^12, 13, 14^.

The anti-inflammatory cytokine IL-10 has been shown to have potent atheroprotective effects in mice^15, 16^. For example, IL-10 deficiency exacerbates atherosclerosis, associated with an increase in plaque lymphocyte and macrophage accumulation^17^. Conversely, intramuscular IL-10 gene transfer reduces plaque area and macrophage infiltration^18, 19^. However, its low molecular weight, rapid clearance, and multiple effects on the immune system have limited its therapeutic potential. To overcome these limitations, engineering approaches are needed to target IL-10 directly to atherosclerotic plaques.

IL-10-encapsulated nanoparticles that target collagen IV^20^ or the αvβ3 integrin receptor^21^ have been developed for atherosclerosis. While these particles have shown efficacy in reducing inflammation in mice, low IL-10 loading capacities (0.5-2.5 wt%) result in the injection of excess foreign polymeric material. Furthermore, these targets are not directly associated with immunogenic molecules or immune cell populations in the plaque. New strategies to target atherosclerotic plaques, for example through antibody fusion proteins, could aid in the clinical translation of IL-10 therapy^22^. Here, we use protein engineering to develop a fusion protein in which IL-10 is fused with an antibody fragment (Fab) that binds to LDL. Upon injection into mice, these Fab-IL-10 constructs bind circulating LDL, accumulate in atherosclerotic plaques, and suppress vascular inflammation (Fig. 1a).

**Figure 1.**
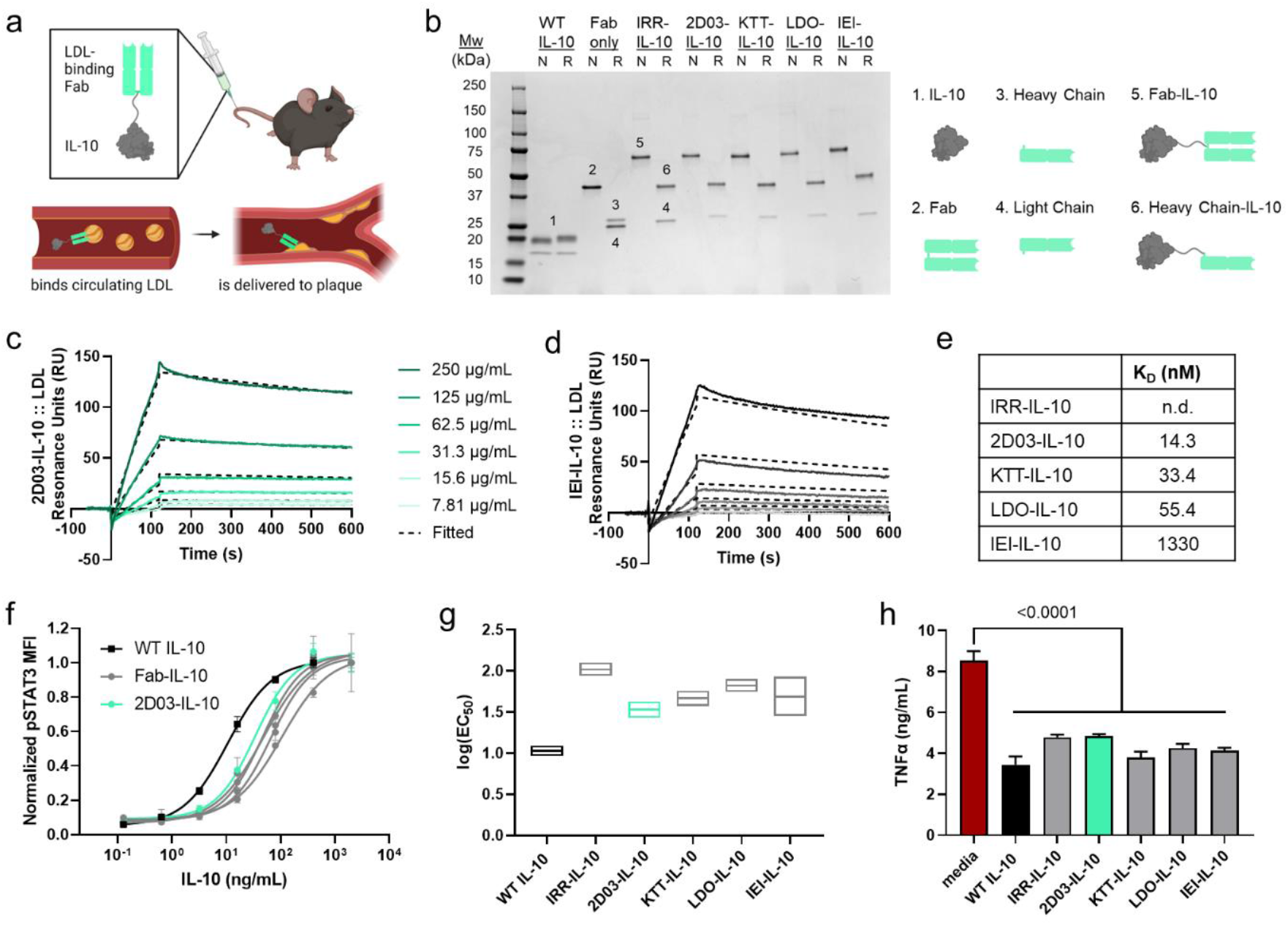
Fusing IL-10 to an LDL-binding antibody fragment enables lipoprotein binding while maintaining signaling and anti-inflammatory properties. a) Conceptual schematic of plaque-targeted IL-10. b) SDS PAGE gel showing various Fab-IL-10 constructs. N, non-reducing conditions. R, reducing conditions. c-e) Binding affinity of Fab-IL-10 to LDL measured using surface plasmon resonance. KD, dissociation constant. Dashed lines represented calculated fit. f) Activity of IL-10 and Fab-IL-10 by phosphorylation of STAT3 (pSTAT3), measured by flow cytometry of RAW 264.7 cells incubated with the indicated concentrations of IL-10 (n = 3). g) Log(EC50) calculated from fitted curves in (f), shown with 95% confidence intervals. h) TNFα secretion of LPS-stimulated RAW 264.7 cells incubated with WT IL-10 or Fab-IL-10 (n = 5). Experiments were performed twice with similar results. Data represent mean +/-standard deviation (f) or mean + standard deviation (h). Statistics performed by one-way ANOVA with Dunnett’s post-test compared to media.

## RESULTS

### Engineered Fab-IL-10 constructs bind to LDL and maintain IL-10 bioactivity

We first identified antibodies from the literature that were developed to neutralize oxidized LDL (oxLDL), a pro-inflammatory lipoprotein associated with atherogenesis^23^, as a passive immunization therapy^24, 25, 26^. Using the complementarity determining regions from four different oxLDL-binding antibodies [2D03, KTT-B8 (KTT), LDO-D4 (LDO), and IEI-E3 (IEI)] and one antibody of irrelevant specificity (IRR) as a size control, we designed plasmids for fusion proteins comprised of antigen-binding fragments bound to IL-10 by a flexible amino acid linker (Fab-IL-10). We recombinantly expressed these proteins and analyzed them by SDS polyacrylamide gel electrophoresis (SDS PAGE; Fig. 1b). Under nonreducing conditions, the full Fab-IL-10 constructs show bands around 70 kDa. Under reducing conditions, they separate into constituent heavy chain-IL-10 and light chain.

We then determined the binding affinity of the Fab-IL-10 constructs to native LDL and oxLDL by surface plasmon resonance (SPR). 2D03-IL-10 shows the highest affinity to native LDL with a dissociation constant (KD) of 14.4 nM (Fig. 1c,e), while IEI-IL-10 has relatively low affinity to native LDL (KD = 1330 nM, Fig. 1d,e). All Fab-IL-10 constructs bind strongly to oxLDL (KD ∼ 10-50 nM, Fig. S1a,c), with the exception of IRR-IL-10, which has no detectable affinity to either species by SPR (Fig. 1e, S1b,c).

After confirming the Fab arm of the construct maintains binding ability, we sought to evaluate whether the IL-10 arm retained bioactivity. Based on STAT3 phosphorylation in a macrophage-like cell line (RAW 264.7), all Fab-IL-10 constructs remained active, albeit with EC50s ∼3 to 10 times less than that of wild type (WT) IL-10 (Fig. 1f,g). Moreover, all Fab-IL-10 constructs reduced TNFα secretion following lipopolysaccharide (LPS) stimulation (Fig. 1h) without affecting cell viability (Fig. S2), indicative of their maintained anti-inflammatory effect.

### LDL-binding Fab-IL-10 constructs exhibit enhanced half-lives in circulation and bioavailability in target tissues

We next evaluated the pharmacokinetics of WT IL-10 and Fab-IL-10 constructs in mice deficient for apolipoprotein E (apoE^-/-^) that were fed a high-fat diet (HFD; 42% from fat) for 10 weeks as a model of atherosclerosis. The small protein WT IL-10 was quickly cleared from the blood, while the larger Fab-IL-10 constructs had longer half-lives (Fig. 2a). After 2 hours, significantly higher concentrations of IL-10 were detected in serum of mice that had received 2D03-IL-10 or KTT-IL-10 compared to WT IL-10 (Fig. 2b). To test if this persistence was due to binding to LDL, we separated very low-density lipoprotein (VLDL) and LDL from the rest of the plasma and measured the amount of IL-10 in the (V)LDL fraction. Supporting the results from SPR, ∼60% of serum IL-10 was bound to (V)LDL for groups receiving 2D03-IL-10, KTT-IL-10, and LDO-IL-10 (Fig. 2c). Mice receiving IRR-IL-10 or IEI-IL-10 had comparably low amounts of IL-10 in the (V)LDL fraction (∼15%, Fig. 2c).

**Figure 2.**
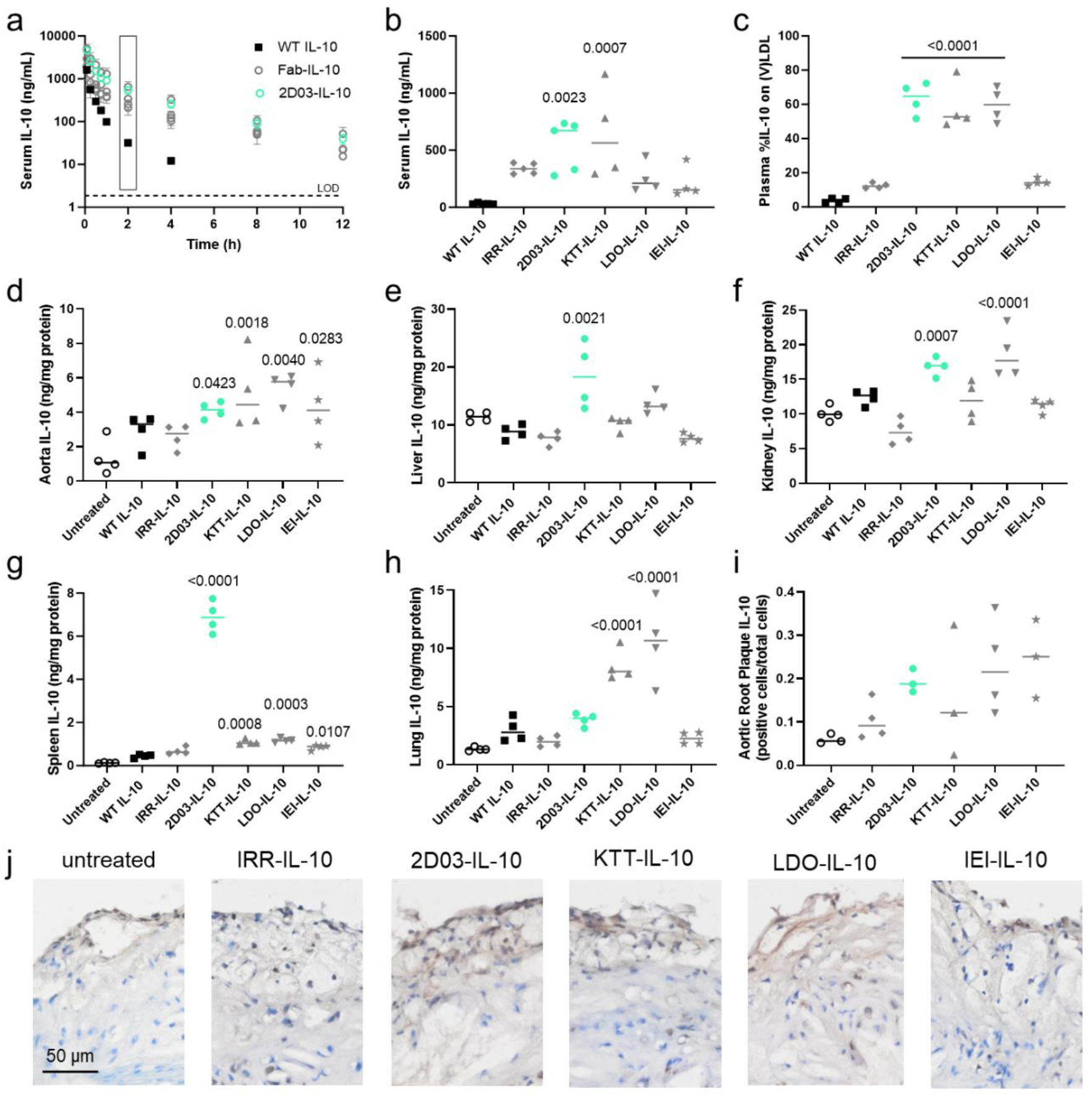
Fab-IL-10 constructs have increased plasma half-lives and tissue bioavailability in apoE^-/-^ mice fed a HFD for 10 weeks. a) Serum concentration of IL-10 following i.v. injection of 10 µg WT IL-10 or equimolar Fab-IL-10. b) Serum concentration of IL-10 in (a) after 2 hr. c) Percent of plasma IL-10 bound to LDL or VLDL in (a) after 2 hr. Concentration of IL-10 in d) aorta, e) liver, f) spleen, g) lung, and h) kidney 2 hr after i.v. injection of 10 µg WT IL-10 or equimolar Fab-IL-10. i) IL-10 in aortic root atherosclerotic plaques by quantified immunohistochemistry. j) Representative images of aortic root with nuclei shown in blue and IL-10 in brown. Scale bar = 50 µm. Data from (a) and (b) are pooled from two independent studies. Data represent mean +/-standard deviation (a). Data points represent individual mice, and line represents median (b-i). Statistics performed by one-way ANOVA with Dunnett’s post-test; p values < 0.05 compared to untreated are shown.

Using the same 2-hour timepoint, we conducted a biodistribution study to determine the tissue-specific bioavailability of each construct. All LDL-binding Fab-IL-10 constructs yielded significantly higher amounts of IL-10 in the aorta compared to untreated mice (Fig. 2d), validating the premise of this targeting approach. 2D03-IL-10 was the only construct that resulted in increased IL-10 in the liver (Fig. 2e), and both 2D03-IL-10 and LDO-IL-10 resulted in increased IL-10 in the kidney (Fig. 2f). Interestingly, while all LDL-binding Fab-IL-10 constructs led to significantly enhanced IL-10 in the spleen, 2D03-IL-10 resulted in concentrations 55 times those of untreated mice on average (Fig. 2g). Finally, KTT-IL-10 and LDO-IL-10 led to increased IL-10 concentrations in lung, whereas 2D03-IL-10 did not (Fig. 2h), indicative of the potential of 2D03 to avoid pulmonary immunosuppression.

To visualize IL-10 in the plaque, we next sectioned the aortic root, where plaques are prevalent in murine atherosclerosis. We used immunohistochemistry to stain aortic root sections for IL-10 and observed an upward trend of IL-10 in the groups of 2D03-IL-10, LDO-IL-10, and IEI-IL-10 from plaque quantification (Fig. 2i). Representative images from these groups also show higher amounts of brown staining, indicative of IL-10 (Fig. 2j, S3). Overall, these results indicate that LDL-binding Fab-IL-10 has a prolonged half-life in circulation and enhanced tissue-specific bioavailability in several organs.

### 2D03-IL-10 reduces vascular inflammation in atherosclerotic mice

With an understanding of the pharmacokinetics and biodistribution of the constructs, we sought to evaluate their efficacy in reducing vascular inflammation in a mouse model of experimental atherosclerosis. ApoE^-/-^ mice were fed a HFD for 8 weeks, followed by 4 weekly intravenous injections of WT IL-10 or Fab-IL-10 constructs while continuing the HFD (Fig. 3a). One week following the final dose, single-cell suspensions of the aortas were analyzed by flow cytometry (Fig. 3b). Mice treated with 2D03-IL-10 had significantly fewer immune cells (CD45^+^) in the aorta than saline-treated mice, comparable to WT C57BL/6 mice fed a normal chow diet (Fig. 3b). None of the other constructs, including WT IL-10, were significantly different than saline. Similar trends were observed with CD11b^+^F4/80^+^ macrophage populations (Fig. 3c). To test whether treatment with 2D03-IL-10 limits leukocyte infiltration into the aorta at earlier disease stages, we repeated the study with mice fed a HFD for a total of 9 weeks. Total immune cells were similarly reduced, as well as CD3^+^ T cells, while CD19^+^ B cells, CD11b^+^F4/80^+^ macrophages, and CD11c^+^MHCII^hi^ dendritic cells trended lower (Fig. 3d-h). Immune cell populations in the spleen remained unchanged following treatment (Fig. S4).

**Figure 3.**
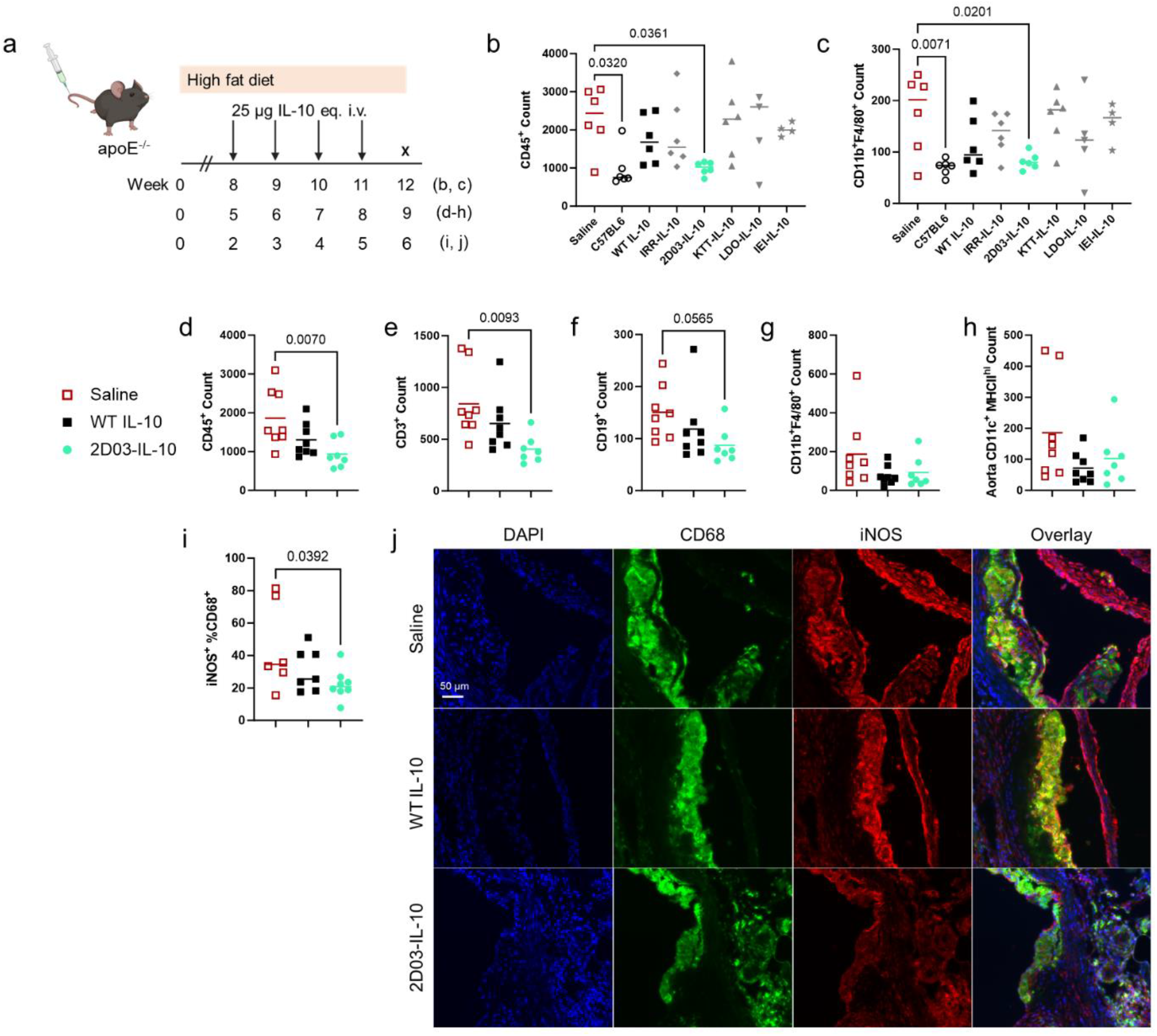
2D03-IL-10 reduces immune cell infiltration into the aorta of apoE^-/-^ mice fed a HFD. a) ApoE^-/-^ mice were fed a HFD for 8 weeks (b, c), 5 weeks (d-h), or 2 weeks (i, j). While continuing the HFD, mice then received 4 weekly i.v. injections of 25 µg IL-10 or molar equivalent Fab-IL-10. Number of b) total immune cells (CD45^+^) and c) macrophages (CD11b^+^F4/80^+^) in the aorta after 12 weeks of HFD by flow cytometry. The C57BL/6 group were aged-matched mice on a normal chow diet. Plots are pooled from n = 3 mice in each of two independent studies. Number of d) total immune cells (CD45^+^), e) macrophages (CD11b^+^F4/80^+^), f) dendritic cells (CD11c^+^MHCII^hi^), g) T cells (CD3^+^), and h) B cells (CD19^+^) in the aorta after 9 weeks of HFD by flow cytometry. i) Quantification of iNOS^+^CD68^+^ cells per total CD68^+^ cells in the aortic root after 6 weeks of HFD. j) Representative images of iNOS staining in the aortic root. Scale bar = 50 µm. Data points represent individual mice, and lines represent median values (b-i). Statistics performed by one-way ANOVA with Dunnett’s post-test; p values < 0.06 compared to saline are shown.

We next analyzed immune cells in the blood one day following the first dose and six days following the final dose of a 9-week study to determine potential effects of 2D03-IL-10 on monocyte recruitment (Fig. S5a). For the first timepoint (one day post-injection), total CD11b^+^ monocytes in the blood remained consistent (Fig. S5b), but the percentage of CCR2^+^ monocytes approximately doubled in mice treated with either IRR-IL-10 or 2D03-IL-10 (Fig. S5c). The CCR2-CCL2 axis mediates monocyte chemotaxis and has been implicated in atherosclerotic inflammation^27, 28, 29^. However, Ly6C^hi^ cells are believed to be the direct precursors to lesional macrophages independent of CCR2 expression^30^, and macrophages alternatively activated by IL-10 are known to have increased expression of CCR2^31^. We found that the percentage of Ly6C^hi^ cells within the population of CCR2^+^ monocytes was similarly reduced by a factor of two in IL-10-treated groups (Fig. S5d, S6), suggesting that this enhanced population of CCR2^+^Ly6C^lo^ cells is anti-inflammatory (Fig. S5e). At the second timepoint (six days post injection), there were no differences in any of these populations between groups (Fig. S5f-h). We also measured plasma cytokines at the later timepoint and found low levels of pro-inflammatory cytokines IL-1β, IL-12, and IL-6, as well as the chemokine CCL2, across all groups (Fig. S7a-e). Plasma IL-10 concentration was significantly higher in 2D03-IL-10-treated mice, even six days following treatment (Fig. S7f). There were no differences between groups in plasma high- or low-density lipoprotein or triglycerides (Fig. S7h), suggesting the reduction in inflammation is independent of cholesterol levels.

To determine the extent of inflammation in the plaque of mice on a HFD for 9 weeks, we then stained sections of the aortic root for lipid content [Oil Red O (ORO)], macrophages/monocytes (CD68), and pro-inflammatory polarization [inducible nitric oxide synthase (iNOS)]. We found that ORO and CD68 staining were consistent throughout the groups (Fig. S8a-d,f). However, 2D03-IL-10-treated mice had significantly lower percentages of iNOS^+^ CD68^+^ macrophages than those treated with WT IL-10 (Fig. S8e). We repeated aortic root staining for a shorter 6-week study and found similar results (Fig. S9), with significantly lower percentages of iNOS^+^CD68^+^ cells in plaques of mice receiving 2D03-IL-10 compared to saline (Fig. 3i,j, S10).

### 2D03-IL-10 targets foamy macrophages

Foamy macrophages are pro-inflammatory lipid-laden cells present throughout all stages of plaque development that have been identified as therapeutic targets in atherosclerosis^32, 33^. Since foamy macrophages are characterized by their accumulation of LDL, we hypothesized that 2D03-IL-10 reduces vascular inflammation by acting on these cells. To test this hypothesis, we isolated splenocytes from atherosclerotic mice and incubated them with 2D03-IL-10, IRR-IL-10, or WT IL-10 ex vivo (Fig. 4a, S11). 2D03-IL-10 showed higher colocalization with all cell types by flow cytometry compared to either IRR-IL-10 or WT IL-10 (Fig. 4b, S12a) and preferentially colocalized with CD11c^+^MHC^hi^ dendritic cells and CD11b^+^F4/80^+^ macrophages in a dose-dependent manner (Fig. 4c, S12b). We found that a higher percentage of macrophages from apoE^-/-^ mice (10 weeks HFD) colocalized with 2D03-IL-10 compared to those from C57BL/6 mice (normal chow diet), whereas colocalization with IRR-IL-10 remained consistent (Fig. 4d).

**Figure 4.**
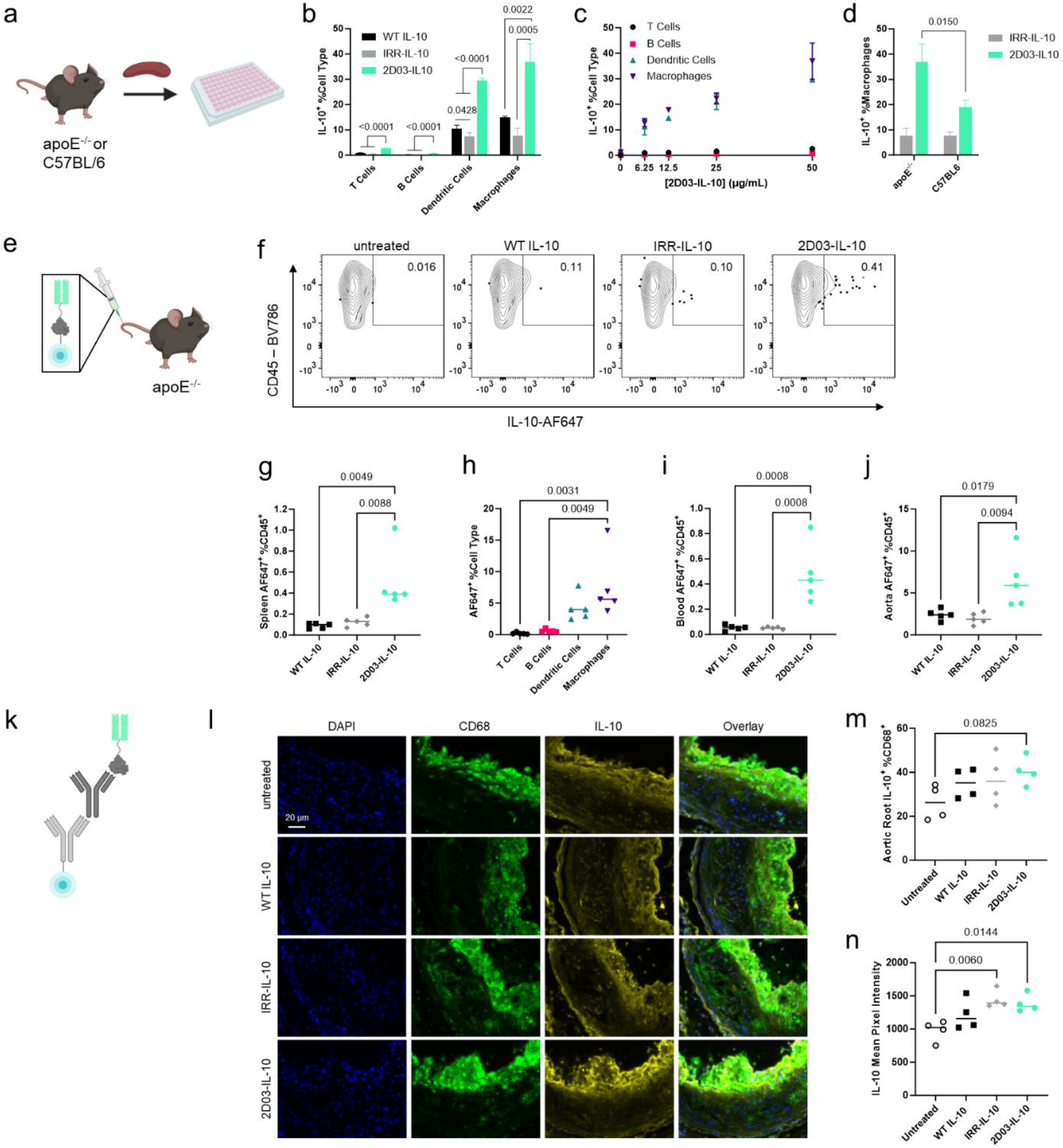
2D03-IL-10 binds immune cells and co-localizes with foamy macrophages. a) Splenocytes from apoE^-/-^ mice fed a HFD for 10 weeks or C57BL/6 mice fed a normal chow diet were plated and incubated with WT IL-10, IRR-IL-10, or 2D03-IL-10 for 30 min at 37 °C. n=3 technical replicates from a single spleen. b) ApoE^-/-^ splenocytes positive for IL-10 by cell type. c) IL-10^+^ apoE^-/-^ splenocytes as a function of 2D03-IL-10 concentration. d) IL-10^+^ macrophages for the different spleen types. e) AF647-labeled WT IL-10, IRR-IL-10, or 2D03-IL-10 was administered to apoE^-/-^ mice fed a HFD for 10 weeks, and organs were harvested for flow cytometry. f) Representative flow cytometry plots of spleens. g) Percent of total CD45^+^ splenocytes positive for AF647. h) Splenocytes positive for AF647 by cell type. i) Percent of total CD45^+^ white blood cells positive AF647. j) Percent of total CD45^+^ aorta-infiltrating immune cells positive AF647. k) IL-10 in the aortic root was detected by immunofluorescence. l) Representative images of CD68 and IL-10 in aortic root plaques. Scale bar = 20 µm. m) Percent of plaque CD68^+^ cells positively stained for IL-10. n) Mean intensity of IL-10 staining in aortic root sections. T Cells: CD3^+^, B Cells: CD19^+^, Dendritic Cells: CD11c^+^MHCII^hi^, Macrophages: CD11b^+^F4/80^+^. The experiments in (a)-(d) were performed twice with similar results. Data represent mean + standard deviation (b,d) or mean +/-standard deviation (c). Data points represent individual mice, and lines represent median values (g-j, m, n). Statistics performed by one-way ANOVA with Tukey’s post-test compared to every other group (a, g-j), two-tailed, unpaired t-test (d), or one-way ANOVA with Dunnett’s post-test compared to untreated (m, n); p values < 0.1 are shown.

We next tested whether AlexaFluor647-labeled 2D03-IL-10 colocalizes with macrophages in an in vivo model (Fig. 4e). Approximately 4 times as many splenocytes associated with 2D03-IL-10-AF647 compared to WT IL-10-AF647 or IRR-IL-10-AF647 by flow cytometry (Fig. 4f,g). Similar to ex vivo data, more splenic macrophages associated with 2D03-IL-10-AF647 than T cells or B cells (Fig. 4h). Additionally, immune cells in the blood and aorta had significantly increased colocalization with 2D03-IL-10-AF647 (Fig. 4i-j). To determine association with macrophages in atherosclerotic plaques, we used immunofluorescence to stain aortic root sections for IL-10 as well as the myeloid marker CD68 (Fig. 4k). We observed a trend toward increased IL-10^+^CD68^+^ cells and significantly higher staining intensity in mice treated with 2D03-IL-10 compared to untreated mice (Fig. 4l-n, S13). Taken together, these data suggest that 2D03-IL-10 associates with immune cells and preferentially colocalizes with foamy macrophages both ex vivo and in vivo.

### Binding LDL is a generalizable strategy

To test whether binding LDL is a generalizable strategy for drug delivery in atherosclerosis, we repeated a subset of studies in LDL receptor-deficient mice (LDLr^-/-^) fed a HFD for 10 weeks (Fig. 5a). Supporting our results in the apoE^-/-^ model, here we found that 2D03-IL-10 has enhanced half-life in circulation and bioavailability in several organs compared to IRR-IL-10 (Fig. 5b,c), suggesting that LDLr does not play an essential role in 2D03-IL-10 trafficking. We also validated in vivo binding to LDL in the plasma of LDLr^-/-^ mice on a HFD, which have lipid profiles more similar to humans^34^, as well as ex vivo binding in plasma of male apoE^-/-^ mice on a HFD and a healthy human donor with normal cholesterol (Fig. 5d-f, S14). Finally, we demonstrate that 2D03-IL-10 broadly binds human aortic plaque cryosections, whereas WT IL-10 shows minimal binding (Fig. 5g). These results demonstrate the generalizability of this strategy to other models and species, indicative of the potential translation of this approach.

**Figure 5.**
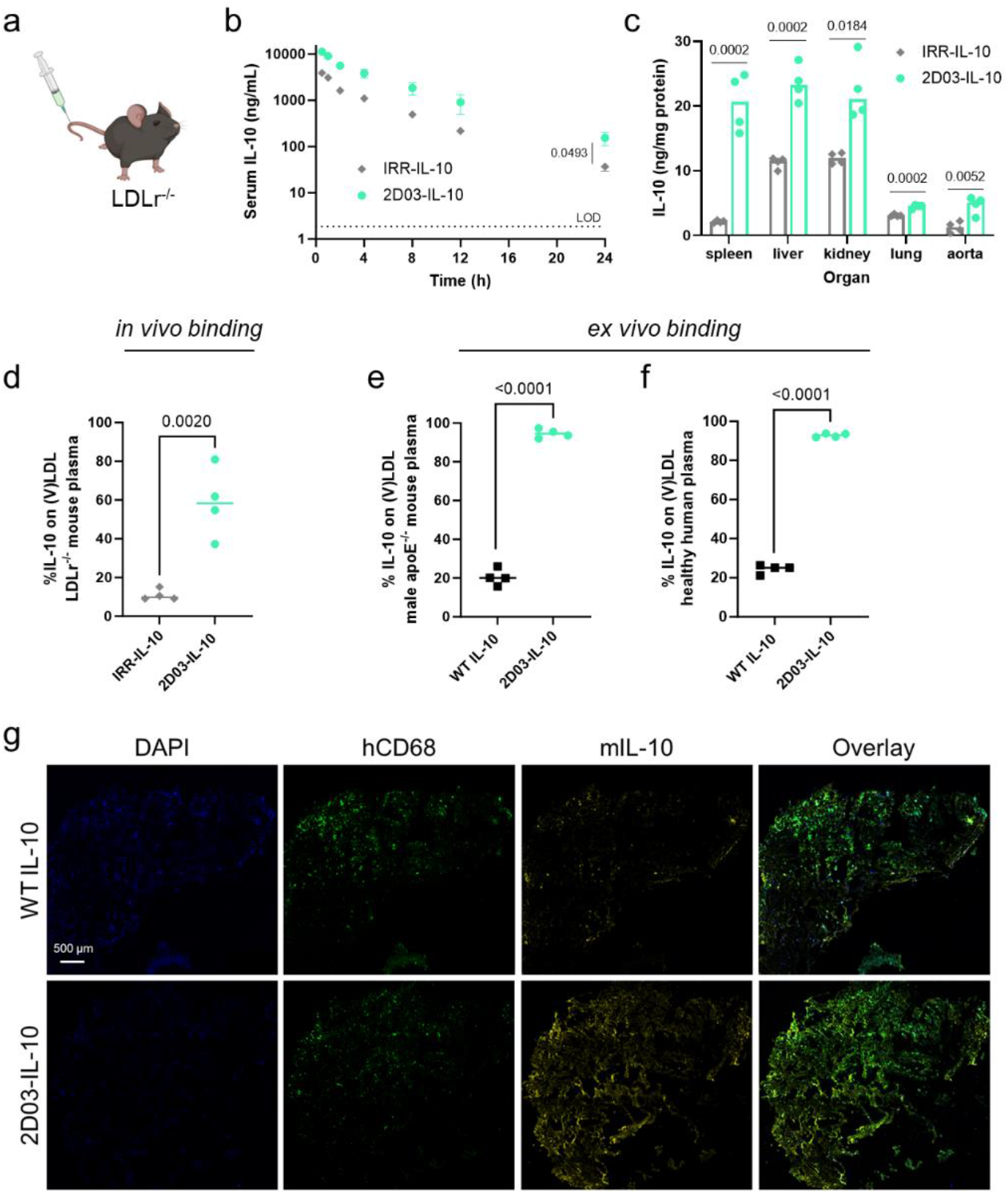
2D03-IL-10 binding to LDL is generalizable to multiple contexts. a) Studies in Figure 2 were repeated in LDLr^-/-^ mice fed a HFD for 10 weeks. b) Serum concentration of IL-10 following i.v. injection of 25 µg (IL-10 basis) IRR-IL-10 or 2D03-IL-10. c) Concentration of IL-10 in spleen, liver, kidney, lung, and aorta 2 hr after i.v. injection. d) Percent of plasma IL-10 bound to LDL or VLDL after 2 hr. e, f) Percent of IL-10 bound to LDL or VLDL upon ex vivo incubation with WT IL-10 or 2D03-IL-10 in male apoE^-/-^ mouse plasma (e) or healthy human plasma (f). g) Binding of WT IL-10 or 2D03-IL-10 to human aortic plaque cryosections. Scale bar = 200 µm. Data represent mean +/-standard deviation (b). Data points represent individual mice, and lines represent median values (c-e). Data points represent technical replicates from one healthy human donor, and line represents median value (f). Statistics performed by two-tailed, unpaired t-test (b-f).

## Discussion

Since oxLDL is a pro-inflammatory species thought to initiate atherogenesis, and naturally occurring antibodies against oxLDL have been identified as atheroprotective^35^, human IgG1 antibodies against oxLDL were recombinantly produced^24^. Specifically, antibodies have been developed against malondialdehyde-modified peptide sequences of apolipoprotein B-100 (apoB-100), the major structural protein of LDL, and used as a passive immunization strategy^24^. Here, we recombinantly expressed antibody fragments from several of these apoB-100-binding antibodies for the purpose of targeting lipid-laden foamy macrophages within atherosclerotic plaques.

We found that Fabs from 3 of 4 of the tested anti-oxLDL antibodies cross-react with human and mouse native LDL, with experimentally determined binding coefficients on the same order of magnitude across the two lipoproteins. Since oxLDL has been found in concentrations on the order of 1000 times lower than native LDL in plasma of coronary heart disease patients^36^, trafficking of the Fab-cytokine fusions is likely dominated by binding to native LDL. In the plaque, oxLDL is in much higher concentrations and may play a larger role in targeting foamy macrophages overexpressing high-affinity scavenger receptors^37^. Future work is needed to better understand the interactions between LDL, oxLDL, 2D03-IL-10, and foamy macrophages.

Interestingly, 2D03-IL-10 has high bioavailability in the spleen, resulting in 55 times higher IL-10 than untreated mice and 6-8 times higher than the other LDL-binding Fab-IL-10 constructs. We hypothesize that the construct binds to or is taken up by circulating monocytes which then traffic to the spleen as a secondary reservoir, as only a small percentage of total monocytes extravasate into the plaque. Since 2D03-IL-10 was the only construct to successfully reduce vascular inflammation, we hypothesize that its ability to bind immune cells and colocalize with foamy macrophages plays a role in its efficacy.

New therapeutic interventions are needed for both early and late stages of atherosclerotic disease. Therefore, we investigated models of experimental atherosclerosis with apoE^-/-^ mice fed a HFD ranging from 6 to 12 weeks, representing fatty streaks in the aorta to widespread lesions. We found that 2D03-IL-10 reduces inflammation across these timescales by decreasing the number of immune cells infiltrating the aorta and reducing pro-inflammatory markers in the aortic root (e.g., iNOS).

In order to validate the results of our studies, we used a second model of atherosclerosis, LDLr^-/-^ mice fed a HFD, with lipid profiles similar to those of humans. While the major lipoproteins in the serum of hyperlipidemic apoE^-/-^ mice are chylomicron and VLDL remnants, LDLr^-/-^ mice accumulate mostly LDL^38^. We observed similar pharmacokinetics and biodistribution of 2D03-IL-10 in this model, suggesting that the approach of targeting LDL is generalizable to different lipid profiles. Additionally, we validated lipid-binding in the plasma of male apoE^-/-^ mice since there are known sex differences in atherosclerotic disease^39^.

Despite evaluating 2D03-IL-10 in two transgenic species, these studies are limited by their use of mouse models of experimental atherosclerosis, which have largely failed to produce results that translate to humans ^34^. However, these Fabs have a human IgG1 framework, and we have shown that 2D03-IL-10 binds LDL in healthy human plasma as well as human aortic atherosclerotic plaques. Additionally, the full 2D03 antibody has a known epitope in human atherosclerotic lesions, which may aid in the translation of this approach^40^. While we use IL-10 as a model cytokine that has been well-studied in atherosclerosis, the modularity of the construct facilitates the targeting of multiple different protein drugs including other cytokines to atherosclerotic lesions. Therefore, we believe this strategy represents a general platform technology for drug delivery in atherosclerosis.

## METHODS

### Fab-IL-10 cloning

The complementarity-determining regions of oxLDL-binding antibodies^26^ were mapped onto a human IgG1 antibody fragment backbone with a lambda light chain, and the sequences were cloned into a human endothelial kidney (HEK) cell expression vector (Abvec2.0). The Fab of irrelevant specificity, which has complementarity-determining regions specific for the xenoantigen outer surface protein A from *Borrelia burgdorferi*^41^, was cloned into the same vector with a kappa light chain, as previously described^42^. DNA encoding murine IL-10 was purchased from Integrated DNA Technologies and cloned into the vectors at the C terminus of the heavy chain following a (GGGS)2 linker. A (His)6 tag was added to the C terminus of IL-10 to enable affinity chromatography.

### Fab-IL-10 expression and purification

HEK293-F suspension cells were cultured in FreeStyle 293 Expression Medium (Gibco) in shake flasks at 135 rpm, 37 °C, and 5% CO2. Cells were transfected at a concentration of 10^6^ cells ml^-1^ with plasmid DNA (1 mg l^−1^) and 25 kDa linear polyethyleneimine (2 mg l^−1^) in OptiPRO SFM (Gibco). After 6 d of culture, the medium was collected by centrifugation and filtered through a 0.22-μm filter. The culture medium was pH-adjusted to 7.4 with Tris HCl, supplemented with 10 mM imidazole, and loaded onto a HisTrap HP 5 ml column (GE Healthcare) at 4 °C using an ÄKTA Pure 25 M (GE Healthcare). The column was washed with 75 mM imidazole [in 20 mM NaH2PO4 and 0.5 M NaCl (pH 7.4)], and protein was eluted at 425 mM imidazole [in 20 mM NaH2PO4 and 0.5 M NaCl (pH 7.4)]. The eluate was then dialyzed against PBS for 24 hr at 4 °C, and protein concentration was determined by measuring absorbance at 280 nm using a NanoDrop (Thermo Scientific). Proteins were verified as >95% pure by SDS-PAGE and contained <0.01 endotoxin units ml^-1^ by HEK-Blue TLR4 reporter assay (InvivoGen).

### Binding to (ox)LDL by SPR in vitro

Surface plasmon resonance measurements were performed using a NTA chip on a Biacore X100 SPR system (GE Healthcare) according to manufacturer’s recommendations. Briefly, Fab-IL-10 (5 mg l^-1^) was immobilized on the chip through His-affinity. LDL or oxLDL from human plasma (Invitrogen) was then flowed over the chip for 60 s in running buffer (PBS supplemented with 0.005% P20 surfactant) at decreasing concentrations. The sensor chip was regenerated with 350 mM EDTA and activated with 0.5 mM NiCl2 for every cycle. Specific binding was determined using the response to a non-functionalized channel as a reference. Binding curves were fitted using BIAevaluation (GE Healthcare) 1:1 binding kinetics to calculate KD.

### STAT3 phosphorylation by flow cytometry

RAW 264.7 cells were cultured in DMEM containing 10% heat-inactivated fetal bovine serum (FBS), 1% penicillin/streptomycin, 1 mM pyruvate, and 50 mM HEPES (all from Gibco) at 37 °C, and 5% CO2. Cells were plated in non-treated 96-well plates (75,000 cells per well) and cultured overnight. The following day, cells were incubated with the indicated concentrations of IL-10 or Fab-IL-10 for 15 min at 37 °C to induce STAT3 phosphorylation. Cells were washed with PBS and detached with dissociation buffer (PBS with 1 mM EDTA and 1 mM EGTA) on ice. Cells were immediately fixed with BD Phosflow Lyse/Fix buffer for 10 min at 37 °C and then permeabilized using BD Phosflow Perm Buffer III for 30 min on ice. Cells were stained with PE-conjugated antibodies against pSTAT3 (pY705, clone 4/P-STAT3, BD Biosciences) at a 1:50 dilution for 1 hr at room temperature in the dark. Data were acquired using a LSR Fortessa flow cytometer (BD Biosciences) and analyzed using FlowJo (v10.8.0, BD Biosciences). The mean pSTAT3 fluorescence intensity of all cells was plotted against cytokine concentration. Dose-response curves were fitted with sigmoidal, 4 parameter logical equations in Prism (v10.1.0, GraphPad) and used to calculate EC50.

### Inflammation inhibition test

To further assess in vitro bioactivity of the Fab-IL-10 constructs, RAW 264.7 cells were plated in non-treated 96-well plates at 10^5^ cells per well. Cells were stimulated with 200 ng ml^-1^ LPS for 12 hr followed by incubation with 100 ng ml^-1^ IL-10 or Fab-IL-10 for 24 hr. Cell supernatants were collected and assayed using a TNFα ELISA (Invitrogen).

### Mice

Female apoE^-/-^ (strain 002052), LDLr^-/-^ (strain 002207), and C57BL/6J mice aged 8 to 10 weeks were purchased from the Jackson Laboratory. After a period of acclimation, mice were switched to a high-fat diet (HFD; 42% from fat, Teklad TD.88137) for the indicated amount of time. All experiments were performed with approval from the Institutional Animal Care and Use Committee of the University of Chicago.

### Pharmacokinetic study

ApoE^-/-^ or LDLr^-/-^ mice that had been on a HFD for 10 weeks received 10 μg (apoE^-/-^) or 25 μg (LDLr^-/-^) equimolar equivalent of Fab-IL-10 i.v. At indicated time points, 6 μl of blood was collected from the tail vein and allowed to clot at room temperature for 30 min. Serum was separated through centrifugation, diluted, and assayed with an IL-10 ELISA (Invitrogen). IL-10 and Fab-IL-10 generated in our laboratories served as the standards for their respective ELISAs. Endogenous IL-10 was below the limit of detection in naive mice.

### Biodistribution of Fab-IL-10 constructs

ApoE^-/-^ or LDLr^-/-^ mice that had been on a HFD for 10 weeks received 10 μg (apoE^-/-^) or 25 μg (LDLr^-/-^) of IL-10 or an equimolar equivalent of Fab-IL-10 i.v. After 2 h, 100 µl of blood was collected for in vivo LDL binding. The heart, aorta, liver, spleen, lung, and kidney were then harvested from mice following perfusion with PBS and euthanasia. Organs were homogenized using a FastPrep tissue homogenizer (MP Bio) in Lysing Matrix D tubes (MP Bio) containing Tissue Protein Extraction buffer (T-PER, Thermo Scientific) supplemented with protease inhibitor tablets (Roche). IL-10 was quantified by ELISA (Invitrogen) and normalized by total protein content from a Pierce BCA Protein Assay (Thermo Scientific).

### Binding to LDL in vivo and ex vivo

For in vivo binding, very low- and low-density lipoprotein [(V)LDL] were separated from the plasma above using a HDL and LDL/VLDL Quantitation Kit (Sigma-Aldrich) according to manufacturer’s instructions. For ex vivo binding, naïve plasma was incubated with WT IL-10 or Fab-IL-10 constructs (500 ng ml^-1^) for 2 hr at room temperature. In both cases, the amount of IL-10 in the HDL-rich fraction and the (V)LDL-rich fraction was quantified with an IL-10 ELISA (Invitrogen) and used to calculate the fraction of bound IL-10. Healthy human plasma was purchased from Lonza Biosciences.

### Sectioning and IHC staining of aortic root

Hearts were fixed in 4% paraformaldehyde for 24 hr and cryopreserved with sucrose (15% sucrose for 24 hr, 30% sucrose for 24 hr, 1:1 30% sucrose and OCT) before being embedded in OCT (Fisher Scientific). Frozen embedded samples were sectioned in 8 µm thickness until all three leaflets of the aortic root were visible. Sections were stained with anti-mouse IL-10 (1:100, Bioss, bs-0698R), followed by ImmPRESS HRP goat anti-rabbit IgG and ImmPACT DAB substrate (Vector Laboratories) according to manufacturer’s instructions. Sections were counterstained with hematoxylin QS (Vector Laboratories). Images were acquired using an Olympus VS200 slide scanner and analyzed using QuPath (v0.5.0).

### Flow cytometry of aorta

Mice were pressure perfused with 30 mL of PBS via the left ventricle after severing the inferior vena cava. The heart with the ascending and descending aorta to the diaphragm was harvested. The perivascular adipose tissue was carefully removed by microdissection while keeping the adventitia intact. The aorta was then severed from the heart at the base, placed in digestion media, and cut into ∼5 mm long pieces. Single cell suspensions were generated as previously described^43^, with minor modifications. Digestion media consisted of DMEM (Gibco) supplemented with 0.5 mM CaCl2 and 2.5 mM MgCl2, 125 U ml^-1^ collagenase XI (Sigma-Aldrich), 450 U ml^-1^ collagenase I (Sigma-Aldrich), 60 U ml^-1^ hyaluronidase, (Sigma-Aldrich), and 60 U ml^-1^ DNase I (Sigma-Aldrich). After 45 min of digestion at 37 °C with orbital shaking, aortas were passed through a 70 μm cell strainer, rinsed, pelleted, and plated into a U-bottom 96-well microplate for staining. Cells were washed in PBS and stained for 15 min on ice with 1:500 Live/Dead Fixable Violet Dye (Invitrogen) and 1:200 anti-mouse CD16/32 (clone 93, BioLegend). Cells were washed in PBS supplemented with 2% FBS (FACS buffer) and stained for 30 min on ice with surface antibodies at a 1:200 dilution in a 1:1 mixture of FACS buffer and Brilliant Stain Buffer (BD Biosciences). Cells were washed with FACS buffer and PBS and fixed with 2% paraformaldehyde on ice for 20 min. Cells were washed twice with PBS and resuspended in FACS buffer. Precision count beads (BioLegend) were added and used to determine absolute cell counts. Data were acquired on LSR Fortessa flow cytometer (BD Biosciences) and analyzed using FlowJo (v10.8.0, BD Biosciences). All antibodies used for flow cytometry are listed in Table S1.

### Binding to splenocytes ex vivo

Single cell suspensions were obtained by passing the spleen through a 70 μm cell strainer. Red blood cells were lysed with ACK Lysing Buffer (Gibco, 3 ml for 5 min). The cells were counted and resuspended in RPMI-1640 medium (Corning) supplemented with 10% FBS and 1% penicillin/streptomycin (Gibco). The cells were plated into a U-bottom 96-well plate (10^5^ cells per well) and incubated with IL-10 or Fab-IL-10 constructs at the indicated concentrations for 30 min at 37 °C or on ice. After three washes with PBS, cells were stained for 15 min on ice with 1:500 Live/Dead Fixable Violet Dye (Invitrogen) and 1:200 anti-mouse CD16/32 (clone 93, BioLegend). Cells were washed and surface-stained with anti-mouse IL-10 (clone JES5-16E3, BD). Cells incubated with IL-10 on ice were then fixed and resuspended as described above. Cells incubated with IL-10 at 37 °C were fixed and permeabilized for 20 min on ice with Cytofix/Cytoperm (BD Biosciences). Cells were washed twice in permeabilization buffer and further stained with anti-mouse IL-10 in permeabilization buffer for 30 min at on ice. Cells were washed twice in permeabilization buffer and resuspended in FACS buffer. Data were acquired on LSR Fortessa flow cytometer (BD Biosciences) and analyzed using FlowJo (v10.8.0, BD Biosciences). All antibodies used for flow cytometry are listed in Table S1.

### Cellular biodistribution study

WT IL-10 and Fab-IL-10 constructs were fluorescently labeled using Alexa Fluor 647 N-hydroxysuccinimide (NHS) ester (Invitrogen), and unreacted dye was removed with a Zeba Spin Desalting Column (7K MWCO, Thermo Scientific) according to the manufacturer’s instruction. WT IL-10 and Fab-IL-10 (5 μg per mouse) were injected intravenously through the tail vein. After 2 hours, the blood was collected, and the spleen and aorta were harvested following perfusion and euthanasia. Red blood cells were lysed following 3 × 3 min incubations with ACK Lysing Buffer (Gibco). The remaining white blood cells were washed with PBS twice and plated for surface staining, as described above. The spleen and aorta were processed and stained as described above.

### Immunofluorescence of aortic root

Frozen aortic root sections were equilibrated to room temperature, rehydrated with phosphate buffered saline (PBS), permeabilized with 10% DMSO, and blocked with 0.05% casein in PBS. Sections were then stained overnight at 4 °C in a humidified chamber with rat monoclonal anti-mouse CD68 (1:400, BioLegend, clone FA-11) and rabbit polyclonal anti-mouse IL-10 (1:25, Bioss, bs-0698R) or rabbit polyclonal anti-mouse iNOS (1:50, Proteintech, 18985-1-AP). Sections were washed twice with PBS supplemented with 0.05% Tween (PBS-T) and then stained for 1 h at room temperature in a humidified chamber with AlexaFluor-488-labeled donkey polyclonal anti-rat IgG (1:500, Invitrogen, A21208) and AlexaFluor-555-labeled donkey polyclonal anti-rabbit IgG (1:500, Invitrogen, A31572). Sections were washed twice with PBS-T and once with PBS before mounting with ProLong Gold Antifade Mountant with DAPI (Invitrogen). Images were acquired using a DMI8 inverted fluorescence microscope (Leica Microsystems) and analyzed using QuPath (v0.5.0).

### Immunofluorescence of human atherosclerosis cryosections

Human cryosections from the aorta of a 76-year-old male with atherosclerosis were purchased from OriGene (CAT#: CS611744). Sections were stained as above with minor alterations. After permeabilization and blocking, sections were incubated with WT IL-10 or 2D03-IL-10 (50 µg ml^-1^ IL-10 basis) at room temperature in a humidified chamber for 2 hr. Washes were performed with PBS supplemented with 0.025% Tween for 1 min each to minimize disruption of plaque lipids. Sections were then stained overnight at 4 °C in a humidified chamber with mouse monoclonal anti-human CD68 (hCD68, 1:400, BioLegend, clone Y1/82A) and rabbit polyclonal anti-mouse IL-10 (mIL-10, 1:25, Bioss, bs-0698R). The following day, sections were stained for 1 h at room temperature in a humidified chamber with AlexaFluor-488-labeled donkey polyclonal anti-mouse IgG (1:500, Invitrogen, A21202) and AlexaFluor-555-labeled donkey polyclonal anti-rabbit IgG (1:500, Invitrogen, A31572). Sections were mounted with ProLong Gold Antifade Mountant with DAPI (Invitrogen). Images were acquired using a DMI8 inverted fluorescence microscope (Leica Microsystems) and formatted using QuPath (v0.5.0).

## Statistical analysis

Statistical analysis between groups was performed using Prism (v10.1.0, GraphPad). Data was assumed to be normally distributed unless otherwise noted. For comparisons of multiple groups, an ordinary one-way ANOVA was used. For comparison of multiple groups to a reference group (e.g., Saline) the ANOVA was followed by Dunnett’s multiple comparisons test. For comparison of multiple groups to every other group, the ANOVA was followed by Tukey’s multiple comparisons test. For comparisons of two groups, a two-tailed, unpaired Students t-test was used.

## DATA AVAILABILITY

The main data supporting the results in this study are available within the paper and its Supplementary Information. Additional processed data are available from the corresponding authors on request. Source data for the figures are provided with this paper.

## ACKNOWLEDGEMENTS

This work was supported by the Chicago Immunoengineering Innovation Center of the University of Chicago, the Gracias Family Foundation, the National Heart, Lung, and Blood Institute T32HL007605-35 (LRV), the American Heart Association Postdoctoral Fellowship Award #916845 (LRV) and the NIH T32 MSTP Training Grants #T32GM150375 and #T32GM007281(SNdM). We thank Catherine Reardon Alulis for helpful conversations regarding murine atherosclerosis models. We thank Suzana Gomes for tissue culture and general laboratory support. We thank the Cytometry and Antibody Technology Core Facility (Cancer Center Support Grant P30CA014599), the Animal Resources Center, the Human Tissue Resource Center, and the Integrated Light Microscopy Core at the University of Chicago. Figures were created with BioRender.com.

## AUTHOR CONTRIBUTIONS

**Lisa Volpatti:** Conceptualization, Methodology, Validation, Formal analysis, Investigation, Data curation, Visualization, Supervision, Project administration, Funding acquisition, Writing – original draft. **Salvador Norton de Matos:** Methodology, Software, Validation, Formal analysis, Investigation, Visualization, Writing – review and editing. **Gustavo Borjas:** Methodology, Software, Formal analysis, Investigation, Writing – review and editing. **Joseph Reda:** Methodology, Investigation, Writing – review and editing. **Elyse Watkins:** Methodology, Investigation, Resources, Writing – review and editing. **Zhengjie Zhou:** Methodology, Investigation, Writing – review and editing. **Mindy Nguyen:** Methodology, Investigation, Writing – review and editing. **Ani Solanki:** Methodology, Investigation, Writing – review and editing. **Yun Fang:** Methodology, Resources, Supervision, Writing – review and editing. **Jeffrey Hubbell:** Conceptualization, Resources, Supervision, Project administration, Funding acquisition, Writing – review and editing.

## COMPETING INTERESTS

The authors declare no competing interests.

## Notes

### Competing Interest Statement

The authors have declared no competing interest.

