## Supplementary Information for "LDL-Binding IL-10 Reduces Vascular Inflammation in Atherosclerotic Mice"

**Table of Contents**

**Supplementary Figure S1.** Binding affinity of Fab-IL-10 to human oxidized LDL (oxLDL).

**Supplementary Figure S2.** Viability of RAW 264.7 cells.

**Supplementary Figure S3.** Lower magnification view of images in Figure 2j.

**Supplementary Figure S4**. Splenic immune cell populations are unchanged upon treatment with 2D03-IL-10.

**Supplementary Figure S5.** IL-10 increases plasma $CCR2^+$ monocytes.

**Supplementary Figure S6**. Gating strategy for flow cytometry of blood.

**Supplementary Figure S7.** Plasma cytokines and cholesterol remain largely unchanged.

**Supplementary Figure S8.** Plaque inflammation, but not lipid content, is reduced upon treatment.

**Supplementary Figure S9.** Plaque area and monocyte content are similar across groups.

**Supplementary Figure S10.** Lower magnification view of images in Figure 3j.

**Supplementary Figure S11.** Gating strategy for flow cytometry of spleens and aorta.

**Supplementary Figure S12.** 2D03-IL-10 binds splenocytes at 4 °C.

**Supplementary Figure S13**. Lower magnification view of images in Figure 4l.

**Supplementary Figure S14**. Cholesterol levels of healthy human plasma.

**Supplementary Table S1.** Probes and antibodies used in flow cytometry.

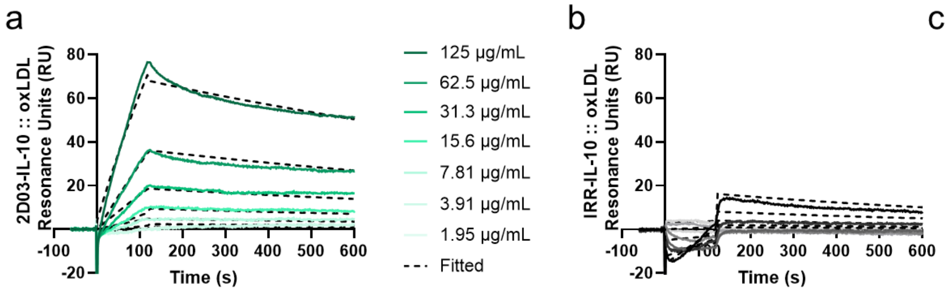


c

| | $K_D$ (nM) |
|---|---|
| IRR-IL-10 | n.d. |
| 2D03-IL-10 | 30.4 |
| KTT-IL-10 | 14.7 |
| LDO-IL-10 | 42.9 |
| IEI-IL-10 | 32.3 |

**Supplementary Figure S1.** Binding affinity of Fab-IL-10 to human oxidized LDL (oxLDL) measured using surface plasmon resonance for a) 2D03-IL-10 and b) IRR-IL-10. c) Summary of dissociation constants ($K_D$), calculated from curves in (a) and (b). Experiments were performed twice with similar results.

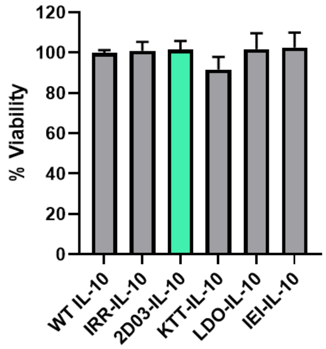


**Supplementary Figure S2.** Viability of RAW 264.7 cells incubated with WT IL-10 or Fab-IL-10 using CellTiter 96 AQueous One Solution Cell Proliferation Assay (MTS, Promega). Percent viability was calculated in comparison to cells incubated with media alone or 1% Triton X. Data represent mean + standard deviation, n=3. Experiments were performed twice with similar results.

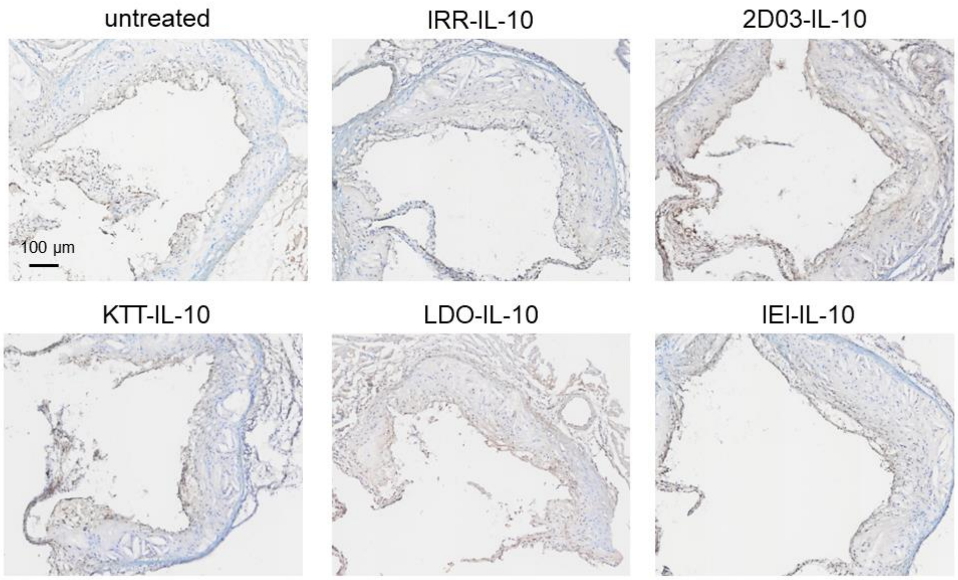


**Supplementary Figure S3.** Lower magnification view of images in Figure 2j, showing one of three leaflets in the aortic root. Scale bar = 100 µm.

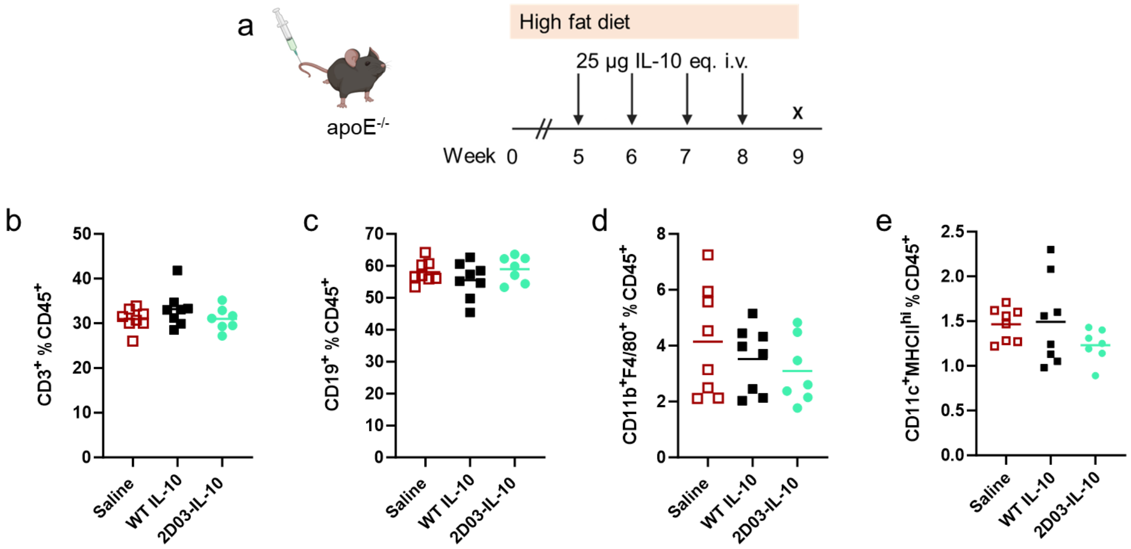


**Supplementary Figure S4**. Splenic immune cell populations are unchanged upon treatment with 2D03-IL-10. a) ApoE$^{-/-}$ mice were fed a HFD for 9 weeks with 4 weekly i.v. injections of 25 µg IL-10 or molar equivalent Fab-IL-10. b) CD3$^{+}$ T cells, c) CD19$^{+}$ B cells, d) CD11b$^{+}$F4/80$^{+}$ macrophages, and e) CD11c$^{+}$MHCII$^{hi}$ dendritic cells as a percent of total immune cells. Lines represent median values. Statistics performed by one-way ANOVA with Dunnett's post-test; no significant differences compared to saline were found.

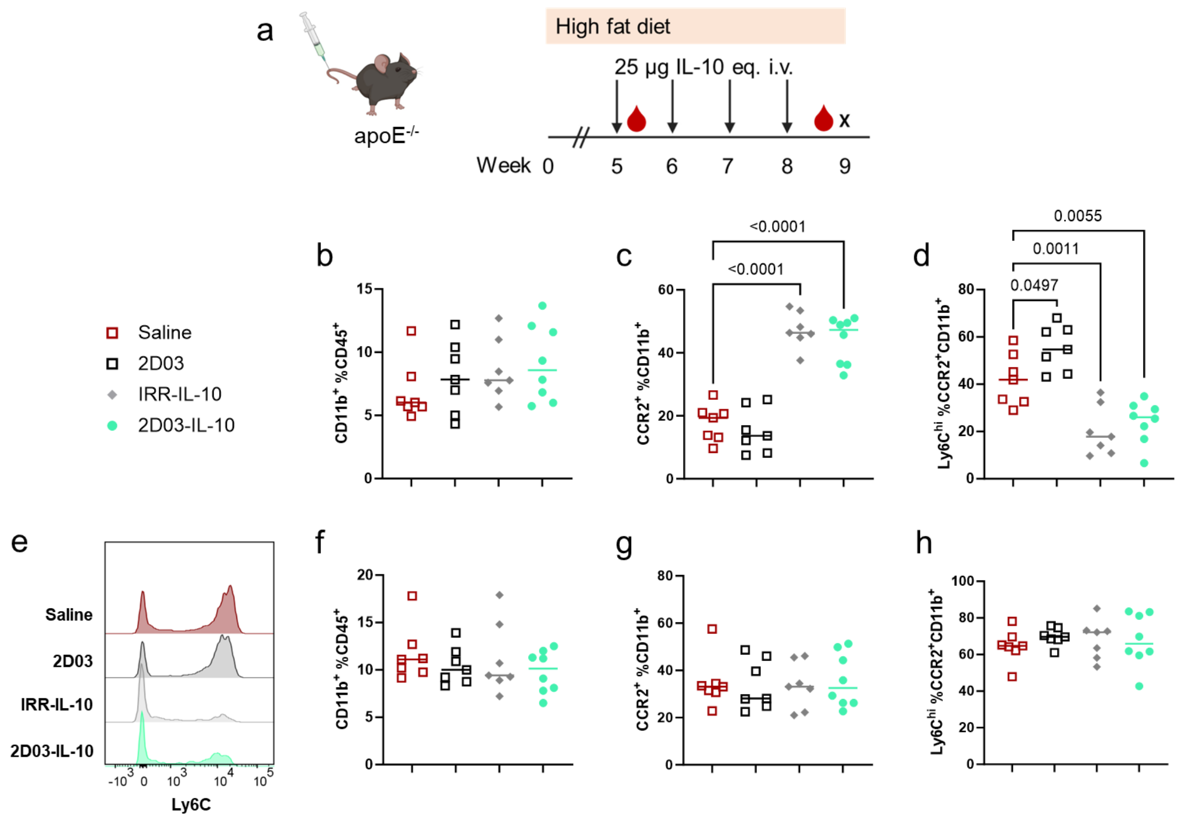


**Supplementary Figure S5.** IL-10 increases plasma CCR2$^+$ monocytes. a) ApoE$^{-/-}$ mice were fed a HFD for 9 weeks with 4 weekly i.v. injections of 25 µg IL-10 or molar equivalent Fab-IL-10. Blood was sampled 1 day after first dose (b-e) or 6 days after final dose (f-j). b, f) Blood monocytes (CD11b$^+$Ly6G$^-$) as a percent of total blood immune cells. c, g) Percent of cells in (b, f) that are CCR2$^+$. d, h) Percent of cells in (c, g) that are Ly6C$^{hi}$. e) Representative flow cytometry histograms of cell populations in (d). Data points represent individual mice, and lines represent median values. Statistics performed by one-way ANOVA with Dunnett's post-test; p values < 0.05 compared to saline are shown.

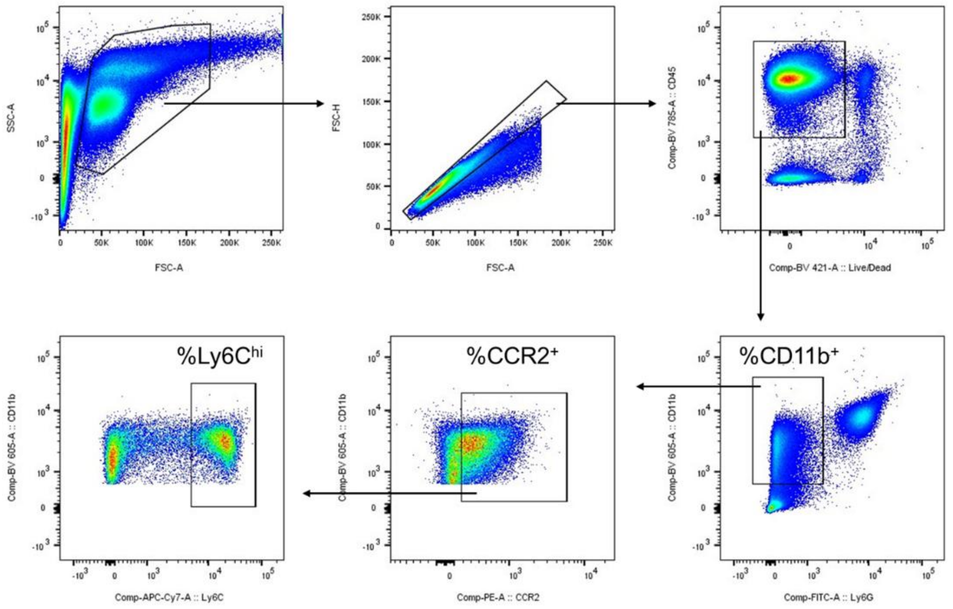


**Supplementary Figure S6**. Gating strategy for flow cytometry of blood.

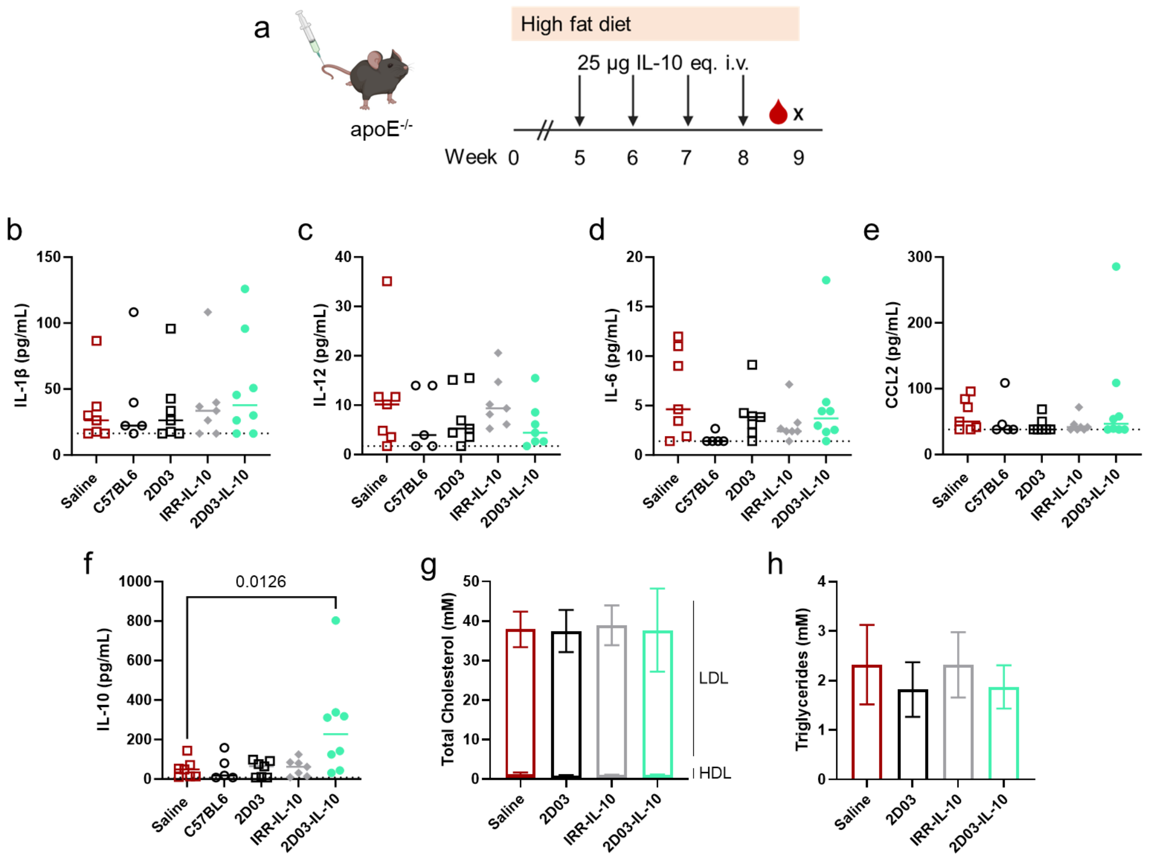


**Supplementary Figure S7.** Plasma cytokines and cholesterol remain largely unchanged. a) ApoE$^{-/-}$ mice were fed a HFD for 9 weeks with 4 weekly i.v. injections of 25 µg IL-10 or molar equivalent Fab-IL-10. Plasma cytokine concentrations of b) IL-1β, c) IL-12, d) IL-6, e) CCL2, and f) IL-10, as measured by a bead-based immunoassay (LEGENDplex, BioLegend). Dashed lines represent assay limit of detection. g) HDL and LDL cholesterol and h) triglyceride levels, as measured with quantification kits (Sigma-Aldrich). Data points represent individual mice, and lines represent median values (b-f). Data represent mean +/- standard deviation, n=7-8 (g, h). Statistics performed by one-way ANOVA with Dunnett's post-test; p values < 0.05 compared to saline are shown.

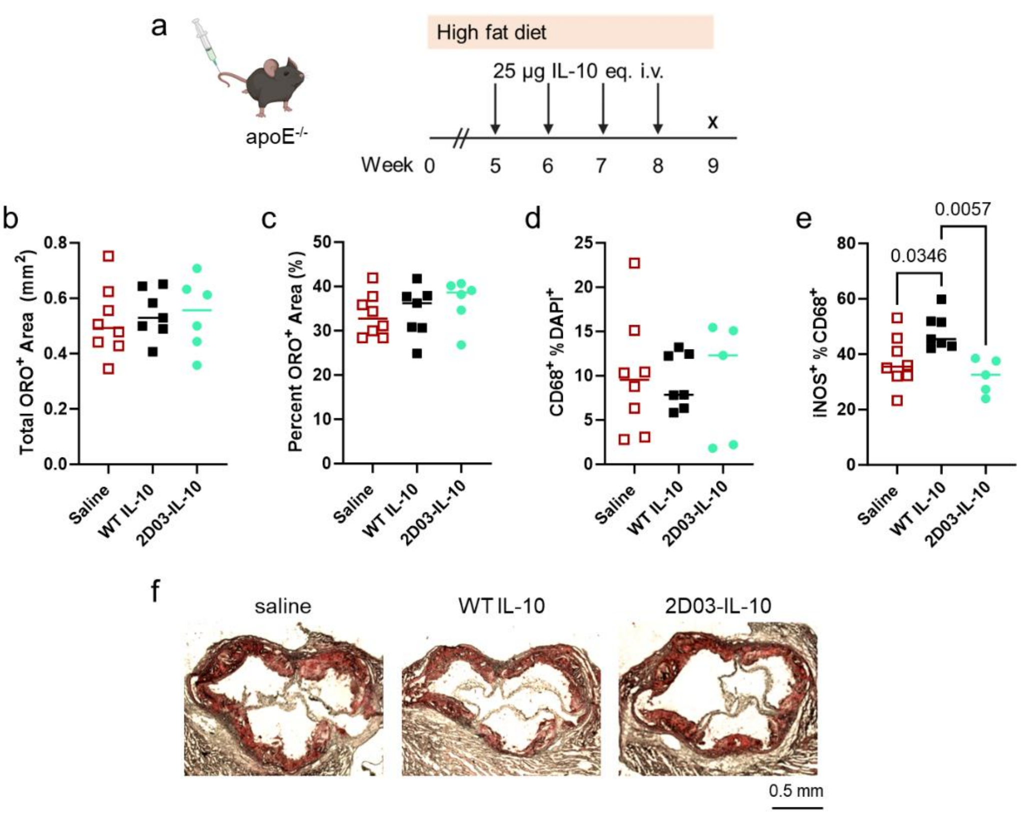


**Supplementary Figure S8.** Plaque inflammation, but not lipid content, is reduced upon treatment. a) ApoE$^{-/-}$ mice were fed a HFD for 9 weeks with 4 weekly i.v. injections of 25 µg IL-10 or molar equivalent Fab-IL-10. Aortic root sections were stained with Oil Red O (ORO, Sigma-Aldrich) and quantified by b) total area and c) percent area. Aortic root sections were stained for d) CD68 and e) iNOS as in Fig. 3j. f) Representative images of ORO staining. Data points represent individual mice, and lines represent median values. Scale bar = 0.5 mm. Statistics performed by one-way ANOVA with Tukey's post-test compared to every other group; p values < 0.05 are shown.

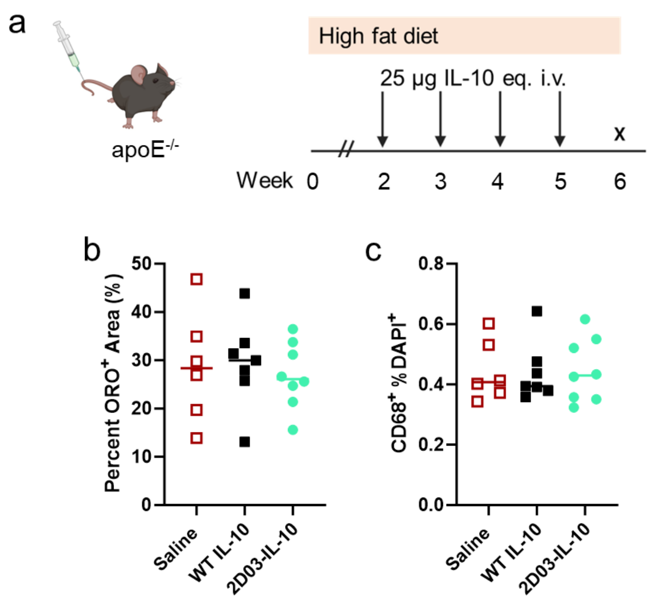


**Supplementary Figure S9.** Plaque area and monocyte/macrophage content are similar across groups. a) ApoE$^{-/-}$ mice were fed a HFD for 6 weeks with 4 weekly i.v. injections of 25 µg IL-10 or molar equivalent Fab-IL-10. Aortic root sections were stained with a) Oil Red O and b) CD68. Data points represent individual mice, and lines represent median values. Statistics performed by one-way ANOVA with Tukey's post-test compared to every other group; no significant differences were found.

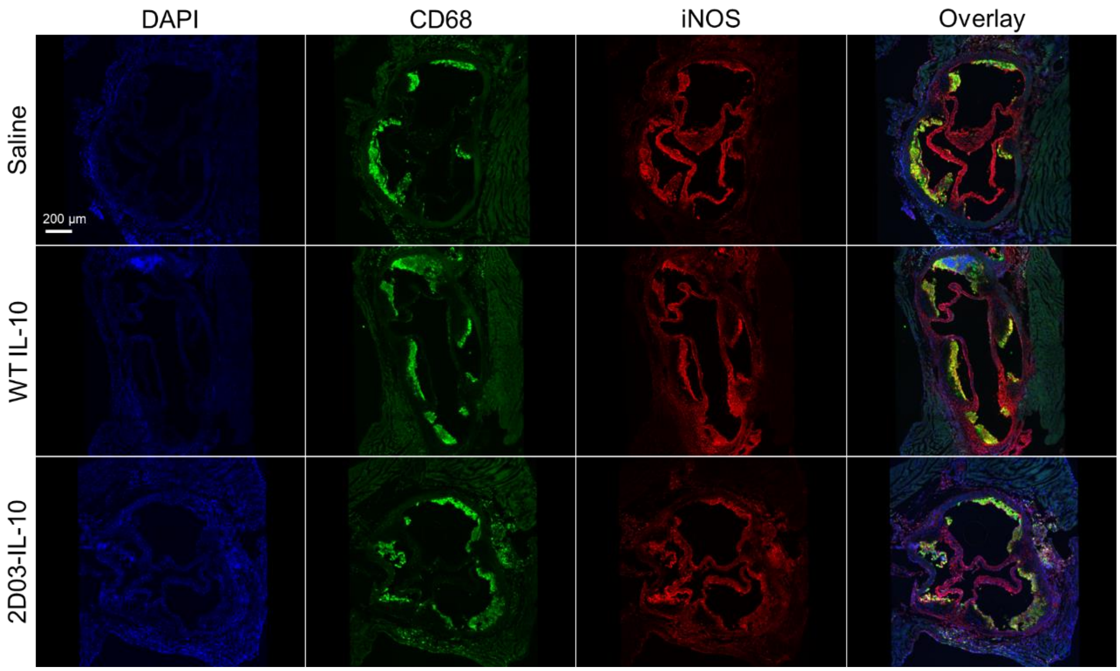


**Supplementary Figure S10.** Lower magnification view of images in Figure 3j, showing all three leaflets in the aortic root. Scale bar = 200 µm.

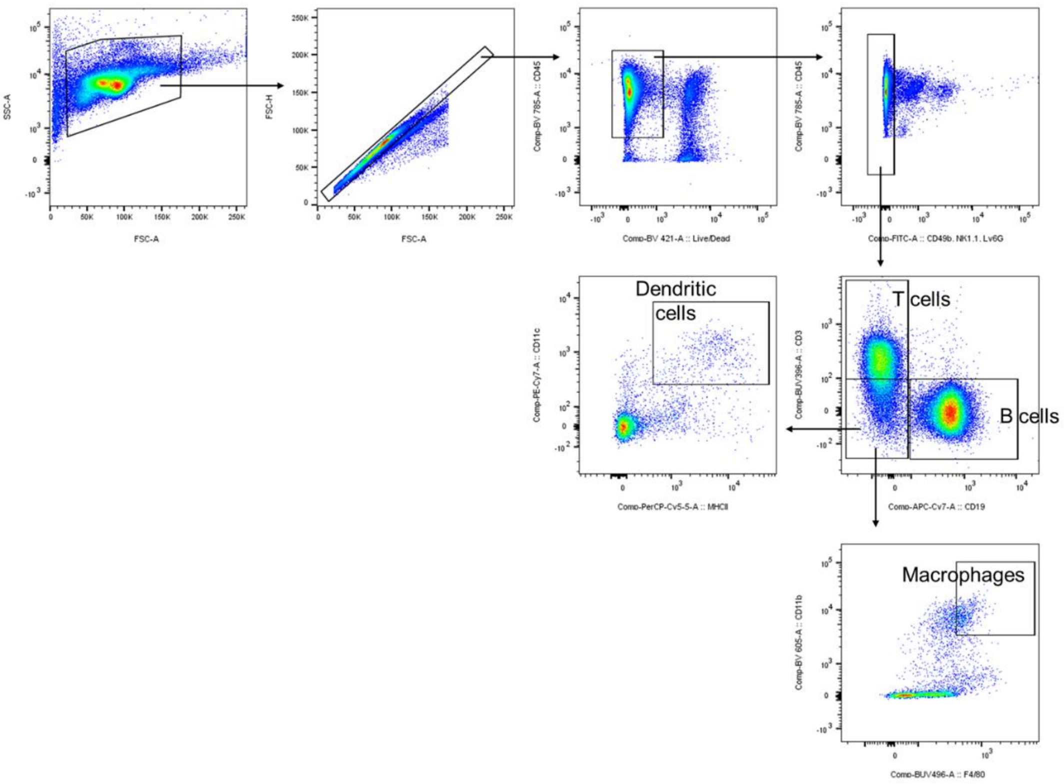


**Supplementary Figure S11.** Gating strategy for flow cytometry of spleens and aorta.

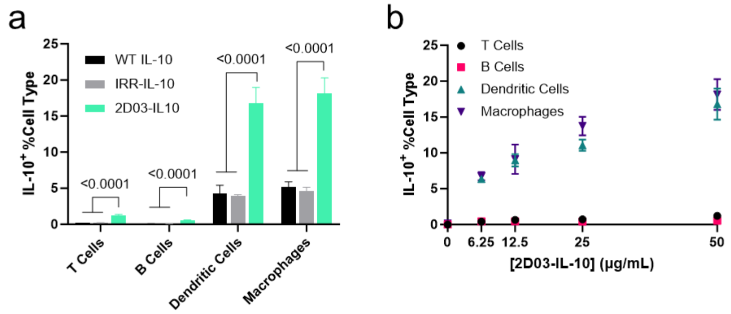


**Supplementary Figure S12.** 2D03-IL-10 binds splenocytes at 4 °C. Splenocytes from apoE$^{-/-}$ mice fed a HFD for 10 weeks were plated and incubated with WT IL-10, IRR-IL-10, or 2D03-IL-10 for 30 min at 4 °C. a) Splenocytes positive for IL-10 by cell type. b) IL-10$^{+}$ cells as a function of 2D03-IL-10 concentration. Experiments were performed twice with similar results. Data represent mean +/- standard deviation. Statistics performed by one-way ANOVA with Tukey's post-test compared to every other group; p values < 0.05 are shown.

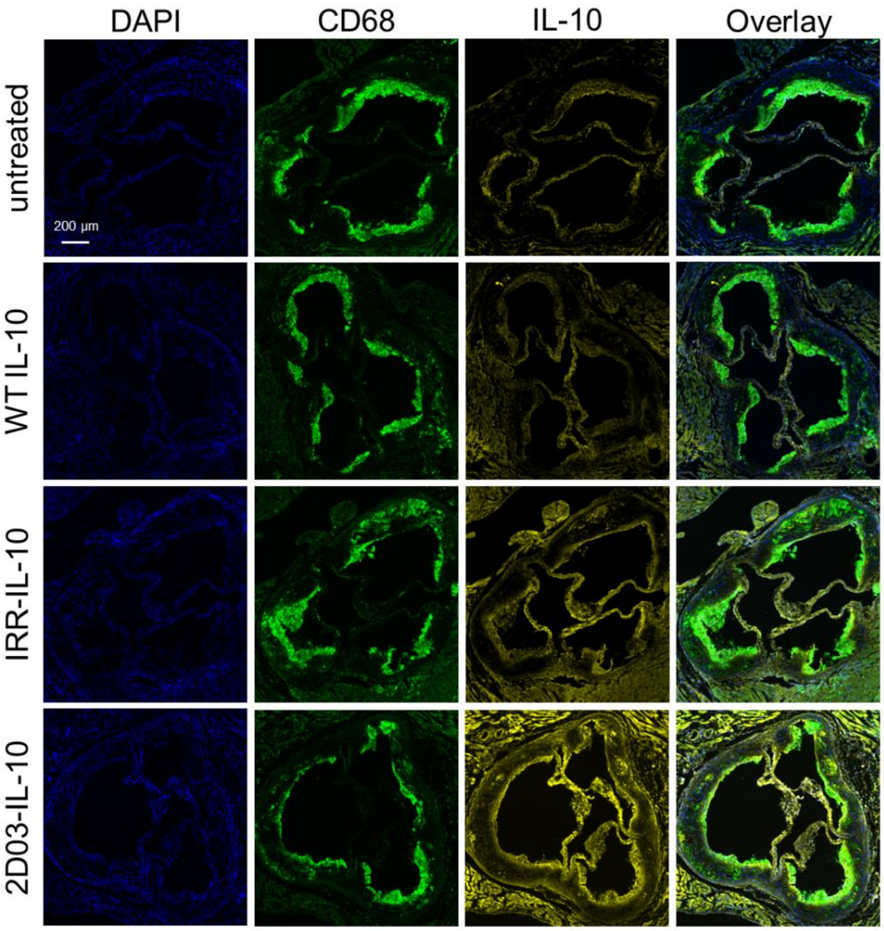


**Supplementary Figure S13**. Lower magnification view of images in Figure 4l, showing all three leaflets in the aortic root. Scale bar = 200 µm.

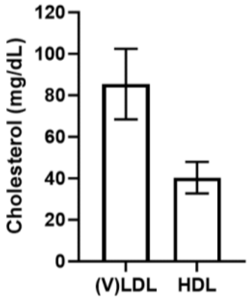


**Supplementary Figure S14**. Cholesterol levels of healthy human plasma. Data represent mean +/- standard deviation; n=4 technical replicates of plasma from one donor.

**Supplementary Table S1.** Probes and antibodies used in flow cytometry.

| Target | Clone | Company |
|---|---|---|
| CCR2 | cat: FAB5538P | R&D Systems |
| CD11b | M1/70 | BioLegend |
| CD11c | HL3 | BD Biosciences |
| CD16/32 | 93 | BioLegend |
| CD19 | 1D3 | BD Biosciences |
| CD3e | 145-2C11 | BD Biosciences |
| CD45 | 30-F11 | BD Biosciences |
| CD49b | DX5 | BD Biosciences |
| F4/80 | T45-2342 | BD Biosciences |
| I-A/I-E (MHCII) | M5/114.15.2 | BioLegend |
| IL-10 | JES5-16E3 | BD Biosciences |
| LIVE/DEAD | (Fixable Violet) | Invitrogen |
| Ly6C | HK1.4 | BioLegend |
| Ly6G | 1AB | eBioscience |
| NK1.1 | PK136 | BioLegend |
